# Ontology-Aware Biomedical Relation Extraction

**DOI:** 10.1101/2022.03.22.485304

**Authors:** Ahmad Aghaebrahimian, Maria Anisimova, Manuel Gil

## Abstract

**Motivation:** Automatically extracting relationships from biomedical texts among multiple sorts of entities is an essential task in biomedical natural language processing with numerous applications, such as drug development or repurposing, precision medicine, and other biomedical tasks requiring knowledge discovery. Current Relation Extraction (RE) systems mostly use one set of features, either as text, or more recently, as graph structures. The state-of-the-art systems often use resource-intensive hence slow algorithms and largely work for a particular type of relationship. However, a simple yet agile system that learns from different sets of features has the advantage of adaptability over different relationship types without an extra burden required for system re-design.

**Results:** We model RE as a classification task and propose a new multi-channel deep neural network designed to process textual and graph structures in separate input channels. We extend a Recurrent Neural Network (RNN) with a Convolutional Neural Network (CNN) to process three sets of features, namely, tokens, types, and graphs. We demonstrate that entity type and ontology graph structure provide better representations than simple token-based representations for RE. We also experiment with various sources of knowledge, including data resources in the Unified Medical Language System (UMLS) to test our hypothesis. Extensive experiments on four well-studied biomedical benchmarks with different relationship types show that our system outperforms earlier ones. Thus, our system has state-of-the-art performance and allows processing millions of full-text scientific articles in a few days on one typical machine.

## Introduction

The job of a biomedical Relation Extraction (RE) system is to identify semantic relationships among biomedical named entities such as genes, drugs, proteins, or chemical substances. There can be a large number of such relationships among different entities. Associations between genes and diseases, interactions among proteins and chemicals, or relationships among drugs and their side effects are a few examples. RE plays an essential role in many biomedical applications such as clinical decision-making or information retrieval. Furthermore, RE is an integral component of Literature-Based Discovery (LBD) systems, commonly used to generate hypotheses for drug repurposing or drug discovery.

The earliest studies in biomedical RE were principally advocating rule-based systems (1–3). Despite being highly precise, rule-based systems are domain-dependent and often require a tedious and labor-intensive procedure for defining the rules.

The advent of modern Machine Learning (ML) paradigms led to a significant boost in the performance of different RE systems, including Chemical-Protein Interactions (CPI) (4), Protein-Protein Interactions (PPI) (5) or Chemical-Induced Diseases (CID) (6) to name a few. Yan et al. (5) use Support Vector Machines (SVMs) (7) for modeling PPI and Onye et al. (6) use SVM and decision trees to model CID. The performance of a typical ML-based system highly depends on the statistical characteristics of the algorithm and the quality of the features.

The process of designing high-quality feature sets known as feature engineering is also a time-consuming process. To address the challenges of feature engineering, several Deep Learning (DL)-based systems with the capacity of self-learning the most meaningful features from complex meaning representations have been developed in recent years.

Many studies on PPI extraction, including Phan et al. (8) and Sun et al. (9) use variants of DL-based algorithms such as Recurrent Neural Network (RNN) (10) or stacked autoencoders. Sänger and Leser (11) and Li et al. (12) employed DL to develop an end-to-end system for adverse drug event and drug-drug relationship detection. Using another DL-based algorithm named Convolutional Neural Network (CNN) (13), Luo et al. (14) proposed segment CNN for RE in clinical notes. Li et al. (15) also made use of RNN to combine the feature vectors trained on MEDLINE with the semantic information obtained from external Knowledge Bases (KB) for relation and entity recognition. However, dependence on an external KB at inference time makes a bottleneck when processing a large volume of data.

Similar to our work, there are a few studies that attempted to integrate different neural architectures. The purpose is to benefit from the advantages and overcome the disadvantages of different shallow and deep algorithms. For instance, Zhang et al. (16) combined RNN and CNN in a hybrid model or Peng et al. (17) combined RNN, CNN, and SVM as an ensemble system.

Textual representations are not the only features that researchers use in their systems. Some studies show the benefits of integrating graph data in RE via leveraging graph data structures. For instance Wang et al. (18) use dependency graph in a Graph Convolutional Network (GCN) (19) to extract Chemical-Disease Relations (CDR). Still, a system capable of using all of these features in a unified model is missing.

Lin et al. (20) and Verga et al. (21) attempted to improve deep sentence-level and document-level RE via the attention mechanism (22). Based on an architecture known as the Transformers (22), Devlin et al. (23) proposed the Bidirectional Encoder Representations from Transformers (BERT). BERT and similar models, including its biomedical variant BioBERT (24) provide a mechanism for generating contextualized language models in contrast to Word2Vec (25) or Glove (26), which generate a static language model.

Contextualized language models in many text analytic tasks, including RE, yield superior results. However, they are considered highly resource-intensive algorithms. Dependence on massive machinery infrastructure usually raises concerns about scalability when considering large-scale RE.

Aiming at developing a large-scale system over the entire full-text articles of PubMed (27), we avoid using any resource-intensive, hybrid, or ensemble system. Instead, we design a unified model that minimizes the load and complexity of the system such that it can process millions of bio fulltext articles in a reasonable time and on a sensible infrastructure.

Our system benefits from the ontology graph and typing information, both enhancing the system’s performance. We train an embedding space for relevant ontology graphs and named entity types at the preprocessing step to ensure that training and inference steps do not depend on an external resource.

We apply our method on four benchmarks with different biomedical relationship types and linguistic characteristics individually to ensure that our model handles agnostic datasets without requiring any particular tuning per dataset. These datasets include ChemProt (28), DDI (29), i2b2 (30), and AGAC (31). Our method shows a substantial improvement (based on F1 score) compared to the current SotA RE systems. In essence, our contributions are as follows:

- We propose a fast multi-channel DL architecture capable of integrating various sources of data into training.
- We introduce type-aware RE and empirically show its impact on the system performance.
- We propose integrating ontology graph data into the model and show that it enhances the results significantly.
- We propose integrating the Unified Medical Language System (UMLS) in the model as a graph-based feature.
- Our system is dataset-agnostic, meaning, although it is trained for each dataset individually, it does not rely on any domain-specific modification for adaptation.

## Methods

Instead of moving towards a more complex DL approach which is less effective (32), we use a simple architecture with several channels that allows us to integrate various sources of data into the training stream without over-complicating the problem.

Meantime ensuring optimum system throughput, to benefit from graph-level and sentence-level information, we train an embedding space on a graph and integrate it into a sentence-level deep neural model. This way, we can enhance the system’s performance while letting it process more than a thousand sentences a second. The required time would be higher by at least one order of magnitude if we would implement it in a graph neural network.

Three sets of features are integrated into our model, namely tokens, entity types, and graph structures extracted from ontologies in the form of graph embeddings. Assume the sentence *S* = *t*_1_, *t*_2_, …, *t*_*n*_ to consist of tokens *t*_*i*_ and to contain two named entities *e*_1_ and *e*_2_. We denote *r* as the relationship pointing to a pair of named entities *e*_1_ and *e*_2_.

In contrast to tokens which are merely occurrences of linguistic units (i.e., words, punctuation marks, symbols, etc.), named entities in life sciences are referred to well-recognized drugs, species, diseases, etc. They may consist of one or more consecutive tokens. Consider the following example:

… of the PDE inhibitors tested, dipyridamole was most effective, with IC50 values of 1.2 and 0.45 microM for inhibition of cAMP and cGMP hydrolysis, respectively.

The named entities are printed in red and blue. For the sake of brevity, we use entity to refer to a named entity from now on. In the ChemProt dataset, *CPR* − 9 is the relationship between the two red entities. Note that there maybe other relationships among the blue entities as well.

The task is then to find *r* such that

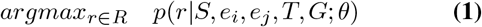

maximizes the probability of *r* where *T* is a set of associated entity types represented in *t* dimensional embedding space, and *G* is a graph consisting of all entities and relations available in the training data. Tokens in *S*, as well as the entities, are represented in *d* dimensional embedding space. *G* also is represented as *g* dimensional embeddings vectors. *R* is a set of relationships, and *θ* are the network parameters. We describe *S, T*, and *G* embeddings in more detail in subsections A, B, and C accordingly.

### A. Token embedding

The most efficient way for representing tokens in almost all NLP tasks is via low-dimensional word vectors, also known as word embeddings. From a broad perspective, word embeddings can be of two types, namely static or dynamic. A static word embeddings algorithm (e.g., Word2Vec (25), Glove (26)) maps each token to a unique low-dimensional vector irrespective of the context where the token occurs. In contrast, a dynamic (i.e., contextual) word embeddings algorithm (e.g., ELMo (33), BERT (23)) maps each token to several different low-dimensional word vectors depending on their surrounding words. Due to the high computational demand of the latter, in this study, we only use static embeddings to ensure a lean and scalable RE system. We use Word2Vec embeddings to represent *S*.

Word2Vec has two variants, namely Continuous Bag of Words (CBOW) and Skip-gram (25). Both variants assume a running window over strings and use a neural network with a single hidden layer for prediction. While CBOW uses the context words in the window to predict the middle word, the Skip-gram uses the middle word to predict the context words. Word embeddings are the network parameters when it is fully trained.

### B. Type embedding

Typing information provides a mechanism for disambiguation when the system is not confident about the relationship between two entities. We integrate type embeddings into the system to examine their impact on the system performance.

In contrast to tokens, there are usually very few types available in a dataset. Consequently, a shallow embeddings technique known as the one-hot encoding (OHE) is sufficient for representing *T*. OHE represents each type as a fixed-length vector where all the values are zero except for the value associated with the type, which is one. The dimension of the one-hot vectors, *t*, is equal to the number of unique types in the relevant dataset.

### C. Ontology graph embeddings

The idea in ontology graph embeddings is to map the graph of an ontology to low-dimensional vectors such that similar components in the graph are close to each other in the low-dimensional space. Therefore, in addition to isolated tokens represented via token embeddings, the network benefits from the information about the interaction of graph components and their neighbors. As the results show in Section G, the embeddings of the ontology graph is a beneficial feature for RE. Graph structures provide three levels of features, namely node, link, and graph as a whole. In the following parts, we describe the features associated with each of these levels.

#### Node-level embeddings

A graph identifies node similarity via different parameters such as having the same neighbors or connecting via the same edge. Either way, similar nodes should be adjacent in their embedding space. Detecting the company that the nodes keep in a graph may improve RE. By node-level embeddings, we provide the network with the positional and structural characteristics of the nodes such as node degree, centrality, and clustering coefficient in a graph.

#### Link-level embeddings

Link-level features are useful for recognizing possible relations between two entities. For a sentence with *N* entities, there are a maximum of *N* (*N* − 1)*/*2 possible relations many of which are not valid. One technique for link validation is to compute a score similar to the PageRank (34) for each node in a pair of nodes (x,y) and get the n highest scored pairs.

#### Graph-level embeddings

Finally graph-level features provide a means of characterizing the structure of the entire graph. When attached to entities and relations, graph-level features might be useful for migrating from one domain to another by projecting the domains into a relevant highdimensional space.

We only estimate and use node-level embeddings to prove the concept and postpone the two other levels to further studies. To set up the input graph for embeddings generation, we construct a graph where entities (i.e., genes, diseases, drugs, etc.) are the vertices, and their relationships are the edges. Transforming this graph into a set of linear random walks (i.e., linearization) is the first step for embeddings generation. After setting the number and the length of random walks, we use a simple sampling agent to linearize the graph. The graph is a directed graph, hence backward moves are not possible. Therefore, at each vertex, the agent decides which outgoing edge to take using a uniform distribution.

Two hyper-parameters, namely the number and the length of random walks, control the agent’s walking behavior. The model uses a portion of training data called the development data to tune these hyper-parameters. After transforming the graph into a set of random walks, we assume each walk as a sequence and use the Skip-gram algorithm to estimate the embeddings.

Due to underlying connections in a graph, splitting a dataset into training and testing sets should be done carefully to ensure that the test set remains unseen. In contrast to textual or imagery data, in graph data, the data points are not isolated from each other instead connected via shared edges. The connections among the vertices of a graph connect data points in training and testing sets either directly or indirectly, compromising the inference by providing the test data with training signals. To make sure that the test set is heldout, we make use of so-called transductive and inductive approaches (35, 36).

In the transductive method, the entire graph is available for all data splits. However, the embeddings algorithm uses only nodes with training labels. In the inductive method, we explicitly remove all shared connections among training, development, and test data such that we have three independent graphs, one for each data split. A disadvantage of this approach is that we remove some beneficial information from the graph.

### D. UMLS graph embeddings

The ontology graph provides a beneficial means of structured data for learning algorithms. However, for some datasets, the ontology graph is not available. A more robust way for generating ontology graph embeddings is to use external resources such as the Unified Medical Language System (UMLS) or Open Biomedical and Biological Ontology (OBO).

We consider the UMLS as an ontology of biomedical concepts. It consists of three main components, namely Metathesaurus, Semantic network, and Specialist lexicon. The Metathesaurus contains over four million biomedical concepts and their associated terms from over 200 source vocabularies. The Semantic network defines 133 broad types (e.g., disease, drug, disorder, etc.) and 54 relationships. It includes semantic types and semantic relationships such as “clinical drug A treats disease B or syndrome C”. Finally, the Specialist lexicon provides lexical information for language processing.

Extracting the clusters of concepts from different vocabularies similar to the UMLS’s Metathesaurus or extracting semantic typing information alike to the UMLS’s Semantic network requires extensive querying among all available ontologies in the OBO Foundry. Given this constraint and for the sake of accessibility and reproducibility, in this study, we use UMLS and postpone OBO integration to further studies.

We extract the words and strings and their associations with their concepts from the UMLS 2021 package. Extracting the concepts, semantic types, and relationships, we construct a semantic graph. After the graph is constructed, a similar mechanism as described in the last subsection projects the concepts and relationships into an embedding space.

### E. Architecture

Recent advances in DL have significantly enhanced RE. Here, we propose a new DL architecture to improve RE over biomedical data (see Figure 1 for the schema). This architecture complements an RNN with a CNN to extract two types of information that are deemed critical in RE. Gated Recurrent Unit (GRU) (37) as an advanced variant of RNNs deals with strings with relatively long dependencies. GRUs in neural networks are often used in form of bidirectional units (i.e., BiGRU). Given a string, one GRU in a Bi-GRU unit extracts the textual features from right to left and the other from left to right. Concatenating the resulting vectors forms a representation that accounts for both directions of the string.

**Fig. 1.**
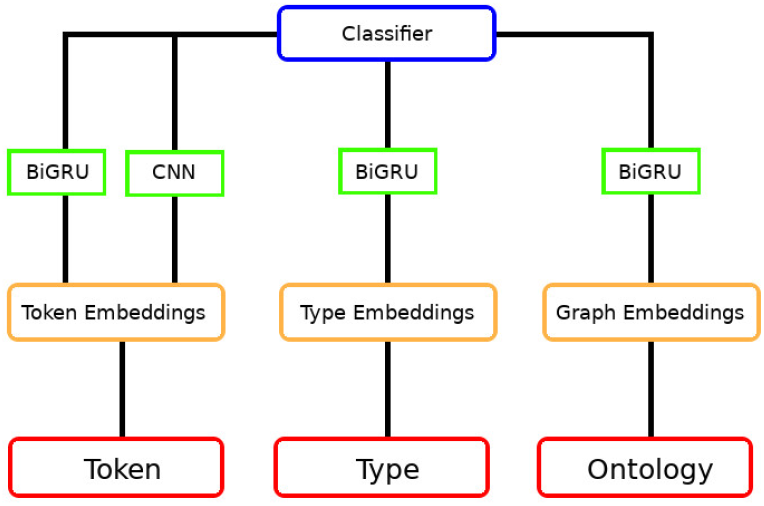
Data-agnostic biomedical RE system architecture.

CNN is a great architecture for keyword or key-phrase extraction. The combination of BiGRU and CNN assures that the model extracts the most informative features with different sequential, local, and time-invariant characteristics. We hypothesize that combining GRU- and CNN-generated features provides RE with a more meaningful representation. Therefore, we propose a Bidirectional Gated Recurrent Unit-Convolutional Neural Network (BiGRU-CNN) multichannel multi-input model for biomedical RE.

This architecture accepts a wide range of features. While tokens and their sequences are valuable features for RE, as we demonstrate via extensive experimentation (please refer to Section H), entity types and ontology graph embeddings facilitate RE as well. Type information helps RE to disambiguate the detected relationships, while ontology embedding provides the model with implicit but beneficial information about entities and their connections in their ontology graph structure.

The first channel in Figure 1 is fed with the isolated token embeddings. While individual tokens provide strong signals for some relationships, the sequence of tokens known as ngrams allows better recognition of some other relationships. The combination of BiGRU and CNN ensures that both of these feature types are extracted. The model concatenates the resulting vectors of BiGRU and CNN to get the overall feature vector. The number of hidden layers for the BiGRU network, sequence length, CNN activation function, the dropout rate, and the optimizer are some of the hyperparameters for this channel.

More recent studies on RE use a form of contextualized word embeddings such as BERT, ELMo, or some of their variants. Computationally, such algorithms are highly demanding with hundreds of millions of parameters. Therefore, to estimate the *S* embeddings in the first channel, we use Word2Vec (Skip-gram) as a static word embeddings algorithm and train it on the PubMed abstracts released by BioASQ (38).

The second channel accepts the type embeddings, and the third channel receives the ontology graph embeddings. The procedure for estimating the embeddings representing *T*, and *G* required for these channels are described in Sections B, and C accordingly. The number and length of random walks for the ontology graph embeddings and the embeddings vector size are two other hyperparameters specific to these channels.

Finally, the classifier on the top is a softmax function which estimates the probability of *r* in Equation 1 by summing up class associated features and normalizing it on the sum of all classes’ features (Equation 2):

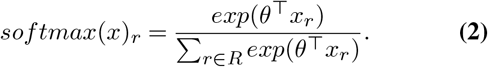

*θ* is the model parameters (as in Equation 1). The variable *x*_*r*_ denotes the features of the relationship *r*. The concatenation of token’s BiGRU/CNN, type’s BiGRU, and ontology’s BiGRU vectors forms the *x*_*r*_ feature vector. The denominator sums over the nominator for all relationships to provide a means of normalization.

The following hyperparameters are reported to ensure reproducibility. All hyperparameters are optimized on the development data if available separately (only for the Chemprot dataset), otherwise on randomly extracted 20% of the training dataset.

## Implementation and Results

### F. Datasets

While many studies in RE focus on a particular dataset, we aim towards designing a dataset-agnostic system. To test the system, we selected four different benchmarks of relationship extraction tasks from various biomedical domains. This selection tries to reflect the thematic diversity, as well as the complexity of the task in terms of sequence length, number of classes, linguistic genre, and vocabulary. We introduce these datasets and some of their statistics in the following subsections.

#### F.1. AGAC

The Active Gene Annotation Corpus (AGAC) is compiled to support drug repurposing. It focuses on the information about functional changes of mutated genes associated with the diseases. The dataset contains 250 and 1000 texts for training and testing sets accordingly. The testing set is not annotated hence the evaluation is performed on the development set (39). The relationship among entities could be one of the following types: gain of function (GOF), loss of function (LOF), too complex to describe (COM), or neutral/unknown (REG) relationships. Table 6 summarizes the statistics of these relationships.

#### F.2. i2b2

The Informatics for Integrating Biology and the Bedside (i2b2) foundation released the i2b2 dataset in 2010. The dataset contains a total number of 394 and 477 annotated training and test samples, respectively. It also includes 877 unannotated reports denoting the relationships among clinical entities (e.g., problem, treatment, and test). Table 2 lists eight classes of relationships included in the dataset.

**Table 1.** System hyper-parameters

| Hyper-parameter | Value | Hyper-parameter | Value |
| --- | --- | --- | --- |
| Emb. size $d$ (tokens) | 200 | Optimizer | adam |
| Emb. $g$ (Ontology) | 128 | hidden layers | 64 |
| Num. random walks | 100 | CNN filters | 32 |
| Length of walks | 16 | CNN kernel size | 4 |
| Drop-out | 0.05 | CNN activation | relu |

**Table 2.** i2b2 classes and the number of supporting samples per train and test sets

| Relationship | Code | training samples | testing samples |
| --- | --- | --- | --- |
| Treatment caused problems | TrCP | 180 | 97 |
| Treatment administered problem | TrAP | 901 | 407 |
| Treatment worsen problem | TrWP | 65 | 10 |
| Treatment improve or cure problem | TrIP | 74 | 26 |
| Treatment not administered due to problem | TrNAP | 62 | 26 |
| Test reveal problem | TeRP | 637 | 264 |
| Test conducted to investigate problem | TeCP | 184 | 83 |
| Problem indicates problem | PIP | 1003 | 452 |

#### F.3. DDI

Drug-Drug Interaction (DDI) (29) dataset is a manually annotated corpus consisting of 792 and 233 texts selected from the DrugBank database and MEDLINE abstracts, respectively. The DDI annotations include 18,502 pharmacological substances and 5028 interactions with an almost perfect inter-rater agreement. Table 7 summarizes the detailed statistics and the types of interactions.

#### F.4. ChemProt

Chemical-Protein Interaction (CPI) recognition plays an essential role in precision medicine and drug discovery in particular, and biomedical research in general (41). The Critical Assessment of Information Extraction (BioCreAtIvE) workshop (28) has several tasks with a focus on protein interactions (42). BioCreative VI challenge provides CHEMical-PROTein interactions (CHEMPROT) dataset for document-level relationship extraction. The corpus contains ten classes of Chemical Protein Relationships (CPR). Nevertheless, only CPR:3-6 and CPR:9 are evaluated in the ChemProt shared task. Chemprot comes with a separate development set (see Table 3).

**Table 3.**
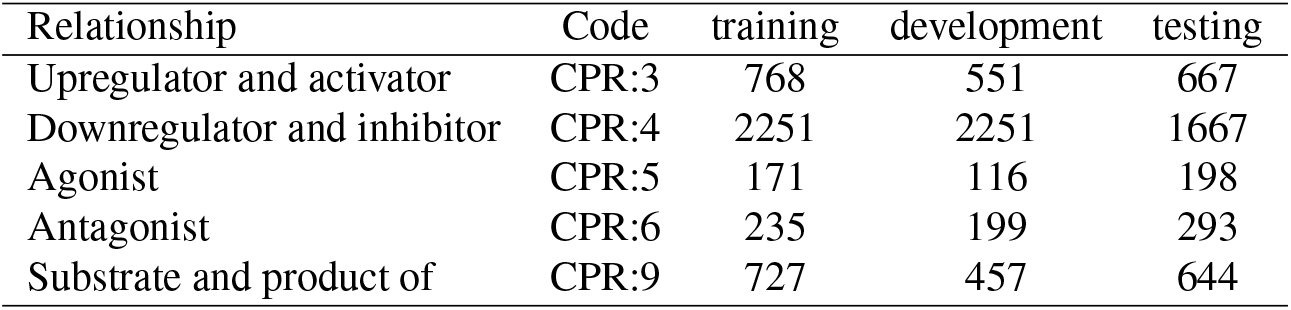
ChemProt classes and the number of supporting samples per train, development and test sets

### G. Results

The following tables report the results of each dataset separately. The hyperparameters are tuned using the grid search strategy. The maximum length of all strings for each dataset is set as the length of the sequences for that dataset.

We use the F1 score as the metric for assessing the system performance to take care of the high imbalance among the number of samples of different classes. The F1 score is computed either per instance (i.e., Micro F1) or per class (i.e., Macro F1). It is the harmonic mean of precision and recall:

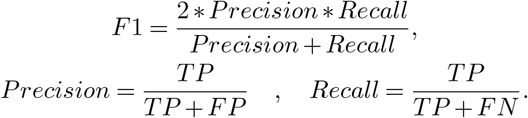

In a multi-class setting *TP* (True Positives) refers to the number of correctly classified samples, while *FP* (False Positives) refers to the number of wrongly classified and *FN* (False Negative) refers to the number of wrongly not-classified samples.

The results in this section are reported based on the ontology graphs generated via the data-driven approach. Although the UMLS-based ontology graphs have a positive impact on the system performance, they yield inferior results compared to the data-driven approach. The distinction between the UMLS-based system and the data-driven approach is reported in the ablation study in Section H. The reason for this inferiority comes from the fact that the coverage rate (i.e., the ratio of entities and relationships in a test set available in the relevant graph embeddings) of the data-driven approach is higher than the UMLS-based approach. Including other biomedical knowledge graphs lead to the model increasing the term coverage hence improving the performance. We postpone this integration to further studies.

A key motivation for our study is to enable to process millions of full-text articles while providing SotA accuracy. Training the models for different datasets takes from less than an hour to at most three hours on a standard machine with an 8-core Core-i7 CPU and 16 GB ram. Depending on the dataset and sequence length of the sentences, the models take a second to make inference over one thousand sentences with an average length of 70 to 120 tokens each. That makes relation extraction for the entire PubMed feasible in a few days and only using one typical machine. Tables 4, 5, 8, and 9 report the results of the system on AGAC, DDI, i2b2, and ChemProt datasets accordingly.

**Table 4.** AGAC test results. The results of the current system are reported with two significant figures. Samples without relationships are extracted as described in Thillaisundaram and Togia (39)

| System | Without none relation |  |  | With none relation |  |  |
| --- | --- | --- | --- | --- | --- | --- |
| Relation/Score | P. (%) | R. (%) | F1 (%) | P. (%) | R. (%) | F1 (%) |
| No-Rel | - | - | - | 95 | 93 | 94 |
| COM | 100 | 100 | 100 | 0 | 0 | 0 |
| GOF | 95 | 82 | 88 | 0.033 | 0.045 | 0.038 |
| LOF | 74 | 87 | 80 | 0.054 | 0.062 | 0.057 |
| REG | 100 | 25 | 40 | 0.031 | 0.042 | 0.035 |
| Current system | **84** | **72** | **78** | **87** | **87** | **87** |
| Thillaisundaram and Togia (39) | - | - | - | 86 | 86 | 86 |

**Table 5.** i2b2 test results. The results of the current system are reported with two significant figures due to the number of test samples. Similar to Table 10 in (40), weighted F-Score is used to ensure a fair comparison.

| System | Yadav et al. (40) |  |  | Current system |  |  |
| --- | --- | --- | --- | --- | --- | --- |
| Relation/Score | P. (%) | R. (%) | F1 (%) | P. (%) | R. (%) | F1 (%) |
| TrCP | 68 | 65 | 66 | 73 | 34 | 47 |
| TrAP | 79 | 82 | 81 | 86 | 94 | 90 |
| TeRP | 87 | 87 | 87 | 83 | 94 | 88 |
| TeCP | 63 | 63 | 63 | 64 | 46 | 54 |
| PIP | 73 | 67 | 70 | 100 | 100 | 100 |
| Macro/Micro score | 74/- | 73/- | 73/- | **81/89** | **74/89** | **76/89** |

**Table 6.** AGAC relationship types and the number of supporting samples per train and development sets

| Relationship | training samples | development samples |
| --- | --- | --- |
| COM | 4 | 1 |
| GOF | 87 | 22 |
| LOF | 93 | 23 |
| REG | 43 | 12 |

**Table 7.** DDI classes and the number of supporting samples per train and test sets

| Relationship | training samples | testing samples |
| --- | --- | --- |
| Advise | 826 | 221 |
| Effect | 1687 | 360 |
| Interaction (Int) | 188 | 96 |
| Mechanism | 1319 | 302 |

**Table 8.** DDI test results. The results of the current system are reported with three significant figures to account for the number of test instances. Similar to (43), the F1 score is micro-averaged F score.

| Relation/Score | P. (%) | R. (%) | F1 (%) |
| --- | --- | --- | --- |
| Advise | 81.9 | 90.0 | 85.8 |
| Effect | 86.0 | 85.3 | 85.6 |
| Int | 94.4 | 35.4 | 51.5 |
| Mechanism | 89.9 | 91.7 | 90.8 |
| Current system | **86.5** | **83.5** | **85.0** |
| Asada et al. (43) | 85.36 | 82.83 | 84.08 |

**Table 9.** ChemProt results.The results of the current and SotA systems are reported in Macro/Micro F1 scores.

| System | Sun et al. (4) |  | Current system |  |
| --- | --- | --- | --- | --- |
| Relation/Score | F1 (%) | P. (%) | R. (%) | F1 (%) |
| CPR:3 | 71.48 | 71.8 | 53.4 | 61.2 |
| CPR:4 | 81.28 | 78.8 | 87.9 | 83.1 |
| CPR:5 | 70.90 | 81.2 | 65.7 | 72.6 |
| CPR:6 | 79.86 | 78.0 | 88.4 | 82.9 |
| CPR:9 | 69.87 | 85.2 | 69.6 | 76.6 |
| Macro/Micro score | - | **79 / 78.8** | **73 / 76.6** | **75.2 / 77.7** |
| Macro/Micro score | 74.6 / 76.5 | - | - | - |

Since there are not enough training data in some classes in the i2b2 dataset, following Yadav et al. (40), we did not use TrWP, TrIP, and TrNAP classes for training and development.

### H. Ablation

This section reports the impact of each layer and several design decisions on the system performance. We limit the parameters of this study to the BiGRU and CNN base models, and the result of adding the type and ontology graph embeddings into the network. The impact of the transductive and inductive approaches for preventing data leakage on ontology graph embeddings are also examined. The ablation study is performed over all datasets to eradicate possible bias as much as possible.

The results in Table 10 show that the base BiGRU configuration consistently outperforms the CNN one, although the performance of the combined model is always higher than the sole BiGRU. It suggests that CNN captures some discriminative features which BiGRU encoders commonly lose.

**Table 10.** Ablation results; the impact of adding each network layer on the system performance. Statistically, significant changes are reported in bold. All scores are reported as Micro F1 score for the sake of consistency.

| Config | Model-Dataset | AGAC(%) | DDI(%) | i2b2(%) | ChemProt(%) |
| --- | --- | --- | --- | --- | --- |
| 1 | Base CNN | 71 | 76.1 | 80 | 70.9 |
| 2 | Base GRU | 72 | 77.2 | 81 | 72.8 |
| 3 | 1 + 2 | 73 | 78.6 | 82 | 74.1 |
| 4 | 3 + Type layer | 75 | 81.4 | 85 | 75.4 |
| 5 | 4 + Ontology layer (UMLS) | 77 | 83.2 | 88 | 77.4 |
| 6 | 4 + Ontology layer (data-driven) | **78** | **85** | **89** | **77.7** |
| 7 | 6 + Inductive training | 77 | 84.5 | 88 | 77.1 |
| 8 | 6 + Transductive training | **78** | **85** | **89** | **77.7** |

Our error analysis empirically shows that CNN does not work well for strictly directional relationships. For instance, CNN makes a lot of mistakes in recognizing CPR:5 and CPR:6 (Agonist and Antagonist relations) in the ChemProt dataset while it recognizes CPR:3 (Upregulator and activator) slightly better than BiGRU.

The impact of type and ontology embeddings layers is also evident from the results. As observed in Table 10, the inductive graph training approach underperforms the transductive one, which is due to the removal of shared connections among different data splits when generating the embeddings. Nevertheless, they both perform better than the models equipped with type embeddings and absolutely better than the base models.

## Discussion

Biomedical relation extraction is a complex task. This complexity is partly due to the linguistic ambiguity and variability inherent in the biomedical entities. The difficulties involved in RE for different linguistic genres such as scientific papers (e.g., ChemProt) versus clinical texts (e.g., i2b2) add to this linguistic complexity. Another reason for such complexity is the wide range of ontologies in life sciences which lead to the definition of numerous relationships’ types. Yet another source of complexity is added to RE because relationships are often directional connections between two entities. However, the text does not always preserve the order of the entities. Similar to the sample in Figure 2 where the second entity occurred before the first one.

**Fig. 2.**
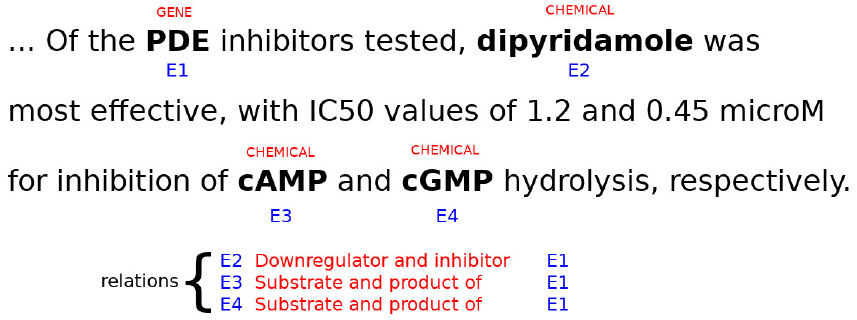
Sample of relationships among different entities.

While addressing the complexity of RE systems requires a comprehensive representation of the model in terms of architecture and the feature sets, the solution should be as fast and scalable as possible such that we can employ it on a large volume of data. It inhibits using overly sophisticated algorithms such as the Transformers (22) which comes with some improved performance at the cost of more demanding computational time and infrastructure.

All studied datasets in this work are highly class-imbalanced that poses a significant issue in multi-class classification tasks. This includes an imbalance among classes as well as an imbalance between positive and negative instances of each class. Class imbalance usually works in favor of the majority class via disregarding the minority class at the training step. TeRP and TrAP in the i2b2 dataset are two evident examples of errors caused by class imbalance. TrCP and TeCP are the worst-performing classes in this dataset; TeCP is often misclassified with TeRP and TrCP is often misclassified with TrAP. In both cases, the class to which the true classes are wrongly assigned belongs to the majority classes.

Another reason for making errors in classification is that in both cases the misclassified classes are semantically similar to true classes; In the first case “Test Conducted to investigate Problem (TeCP)” and “Test Reveal Problem (TeRP)” and in the second case “Treatment Cause problems (TrCP)” and “Treatment Administered Problem (TrAP)” are considerably similar. Other confused classes compromise less severe misclassification. Our experiments on various embeddings show that an embedding trained on biomedical data yields fewer misclassified instances of this type.

The worst-performing class in the DDI dataset is also the minority class *Int* which often is overshadowed by *Effect*. One reason for this is that *Int* is the super-class denoting any interaction which conveys the same semantics as *Effect* may do. DDI also shows fewer misclassified instances when the model uses an embedding trained on biomedical data.

The impact of class imbalance on the system performance is again evident in the ChemProt dataset. CPR:4 as the majority class is a customary target for other classes such as CPR:3 to be misclassified.

Class imbalance is a big problem in all the datasets. One way to rectify this problem is to over-sample the minority classes via textual augmentation methods such as paraphrasing or summarizing models. Another way is to integrate weighted loss which penalizes misclassification on minority classes with more severity. We intend to improve the system following these ideas in future studies.

## Conclusion

Relation Extraction is a fundamental task in biomedical text analytics. There is a wide range of domains within biomedical and health sciences. Therefore a universal model capable of extracting relationships across various biomedical subdomains is highly desirable since it reduces the time and effort required to design domain-specific architectures. Employing graph ontology and biomedical types represented as embeddings, we designed a deep neural network for relation extraction adaptable to various domains given the ontology and type information encoded as an embeddings layer. The network takes this information directly from the datasets in a data-driven approach or indirectly from the UMLS as an external resource. Our system obtains state-of-the-art results on four datasets from different biomedical sub-domains, namely; Chemical Protein Interactions (CPI), Drug-Drug Interactions (DDI), Gene functions, and clinical problems and tests. Due to its uncomplicated yet quick encoders and classifier, it makes relation extraction feasible on a large volume of textual data and within a limited time.

